# Closed-form expressions for the directions of maximum modulation depth in temporal interference electrical brain stimulation

**DOI:** 10.1101/2025.05.16.654079

**Authors:** Mariano Fernández-Corazza, Sergei Turovets, Carlos H. Muravchik

## Abstract

In temporal interference (TI) electrical brain stimulation (EBS), an emerging neuromodulation technique, the interference of two high-frequency currents with a small frequency difference is used to target specific brain regions with better focality than in standard EBS. While the magnitude of the modulation depth has been previously investigated, an explicit formula for the directions in which this modulation is maximized has been lacking. This work provides a novel closed-form analytical expression for the orientation of maximum modulation depth in TI EBS. We also found a secondary orientation where the modulation depth has a local maximum and provide closed-form analytical formulas for this orientation as well as for the modulation depth along this orientation. To our knowledge, this is the first published closed-form expressions for these specific orientations in the context of TI stimulation. We mathematically derive these compact formulas and validate them through comprehensive computational simulations using a realistic human head model. Our simulations demonstrate that the modulation depth predicted with our new analytic direction formula of maximum modulation depth is indeed the maximum compared to other directions. The derived closed-form expression offers a faster and more precise alternative to numerical iterative optimization methods used in previous studies to estimate this direction. We also provide a complete analytical derivation of the widely used (Grossman et al., 2017) formula for the maximum modulation depth magnitude. Furthermore, we found that due to interference in 3D, the modulation depth along the secondary maximum orientation can be of similar strength than the maximum modulation depth intensity when interfering electric field vectors are significantly misaligned. Finally, we show that by modifying the ratio of the injected current strengths, it is possible to steer these directions to fine tune the stimulation along a desired direction of interest. Overall, this work represents a detailed treatment of TI electrical fields in 3D. The presented closed-form expressions for the directions of maximum and secondary maximum modulation depths are relevant for the better interpretation of both simulated and experimental results in TI studies by allowing comparison with neuronal orientations in the brain.

## 1 Introduction

Electrical brain stimulation (EBS) is an emerging therapy for the treatment of neuropsychiatric conditions such as clinical depression (Kalu et al., 2012), Parkinson’s disease (Boggio et al., 2006), anxiety and chronic pain (Mori et al., 2010). Research has also shown that EBS can be a valuable therapeutic tool in epilepsy (Holmes et al., 2019; Yook et al., 2011), stroke rehabilitation (Schlaug et al., 2008), and other neurological and psychiatric disorders (Brunoni et al., 2013). It has also been extensively studied in the context of enhancing cognition including memory and learning (Berryhill and Jones, 2012; Nitsche et al., 2003) and psychomotor skills in military and sports contexts (Perrey, 2023; van der Groen et al., 2025).

This technique may eventually become an alternative to psychoactive drugs, as it can be more selective than drugs by targeting specific regions of interest in the brain with minimal adverse side effects. If applied with electrodes placed in the scalp, EBS is termed Transcranial Electrical Stimulation (TES) or Transcranial Brain Stimulation (TBS). Even without producing direct neuronal firing, the application of TES is capable of modifying cortical excitability (Nitsche and Paulus, 2000; Priori et al., 1998) as well as brain rhythms and networks (Lang et al., 2005; Priori, 2003), which is why the method is also termed Transcranial Electrical Neuromodulation (TEN). If direct or alternating currents are used, TES is termed transcranial direct current stimulation (tDCS) or transcranial alternating current stimulation (tACS), respectively. Despite recent advances, ongoing debates address the clinical effectiveness of TES (Antal et al., 2015; Horvath et al., 2015, 2014), with many issues still to be resolved, in particular, substantial inter-subject response variability (Batsikadze et al., 2013; Wiethoff et al., 2014). The general idea is that optimal targeting protocols and the use of subject-specific accurate head models may enhance rigor and reproducibility in TES (Bikson et al., 2018).

Following the work by (Grossman et al., 2017), there has been a renewed interest in temporal interference (TI) TES introduced clinically in the 1960s in the context of electrosleep (Sachkov et al., 1967). In TI, two current injection patterns are simultaneously active, each of them injecting a relatively high frequency (in a few KHz range) but with a low frequency difference between them (for instance 10Hz). Each of these current injection pairs produces a different electric field *E*_1_ and *E*_2_ with the idea that neither peripheral nerves nor the brain are responsive to each of the signals separately due to its high frequency. However, the neurons of the brain mix input signals nonlinearly and rectify the low frequency differential beating signal. The modulation depth of the interfering fields, which is the amplitude of the low-frequency envelope and thus also called the beating amplitude, is thought to be responsible for the neuromodulation effects (Grossman et al., 2017). Although the field is still in the early stage of systematic experimental research, the recent TI simulations in realistic human head models have shown that neuromodulation with TI is feasible despite being attenuated by the skull like TES. In contrast to ordinary TES, it was demonstrated that TI TES can achieve increased focality and local “hot” spots in deep regions of the brain which is usually impossible in TES (even in optimal TES) where intensity of stimulation diffusively decays away from electrodes (Fernandez-Corazza et al., 2019). Focality gets even better if TI stimulation montages are optimized (Esmaeilpour et al., 2021; Howell and McIntyre, 2020; Huang et al., 2020; Lee et al., 2020). Thus, TI (and optimized TI) offers a very promising approach for specific manipulation of brain networks.

Recent studies have explored the efficacy of TI stimulation in humans (Zhu and Yin, 2023). One study investigated the effects of 20 Hz TI stimulation targeting the primary motor cortex (M1) in 40 healthy individuals (Zhu et al., 2022). Their findings indicated an enhancement in functional connectivity between M1 and secondary motor cortices during stimulation. Similarly, another work examined the impact of TI stimulation on motor cortex excitability and motor learning (Ma et al., 2022). They reported that 70 Hz TI stimulation improved reaction times in a random reaction time task and increased motor cortex excitability, while 20 Hz TI stimulation significantly enhanced motor learning in a serial reaction time task. A recent study provided further evidence of TI’s potential by demonstrating non-invasive stimulation of the human hippocampus, revealing changes in memory-related functional connectivity and behavioral improvements in spatial memory tasks (Violante et al., 2023). Their study supports the feasibility of TI for targeting deep brain regions involved in cognition. Another study combined transcranial TI with virtual reality (VR) and functional magnetic resonance imaging (fMRI) to enhance spatial memory in healthy participants (Beanato et al., 2024). Participants received TI targeting the hippocampus and adjacent structures while navigating VR environments designed to test spatial memory. The study demonstrated that TI, combined with VR training, led to significant improvements in participants′ ability to recall locations and navigate effectively, suggesting a potential non-invasive intervention for age-related memory decline and early-stage dementia. Despite these promising outcomes, challenges persist in the application of TI stimulation in human studies. One significant concern is the limited amplitudes of stimulation currents permissible in humans compared with animal models, which may affect the efficacy of the intervention. Additionally, inconsistencies in neurophysiological responses across studies highlight the need for further research to understand the underlying spatiotemporal distribution of the electric fields, optimize stimulation parameters, and fully understand the underlying mechanisms of TI stimulation in humans.

The spatiotemporal patterns of TI are complex and understanding them is key to developing advanced TI tools and to better interpret the experimental results. At each point in space, TI of two sinusoidal currents with frequencies *f*_1_ and *f*_2_ produces a vector that rotates in time with a carrier frequency (*f*_1_ + *f*_2_)/2 and an amplitude varying with frequency (*f*_1_ − *f*_2_)/2 within a plane defined by the two interfering electric field vectors, as it is pointed in our previous work (Fernández-Corazza et al., 2019) and later in (Huang, 2023). If this rotating vector is projected onto a specific predefined direction of interest, such as the predominant axonal orientation, the formula to obtain the modulation depth is straightforward (Grossman et al., 2017). The problem with rotation is that a dosage delivered in one predefined direction, say, parallel or perpendicular to the cortex, might be very low even with a significant total magnitude of the electric field. However, at each point in the brain, there exists a specific direction among all possible orientations in which beating signal depth is maximized, resulting in the highest dosage. This is why an analytical expression for finding this direction is important. The work by (Grossman et al., 2017) provides a conditional formula for the expression of the maximum intermodulation depth intensity, but it does not provide an expression for the specific direction where this maximum intermodulation depth occurs, nor the derivation of the formula. Several other articles simply replicate this formula without any demonstration of its origin (Rampersad et al., 2019; von Conta et al., 2021). (Wang et al., 2022) simulated the electric fields at a macroscopic and neuronal level, and they provide a different expression for the maximum amplitude depth, which looks similar to that of (Grossman et al., 2017), but we show here that they do not exactly match. (Huang and Parra, 2019) opted not to use (Grossman et al., 2017) formulation, reverting to a numerical solution of the maximization-minimization problem by performing a 1D brute-force search on the angle of the direction of maximum modulation depth. (Lee et al., 2020) cite this study of (Huang and Parra, 2019) and use the same maximization-minimization formulation, which is less efficient than (Grossman et al., 2017) formulation. (Meng et al., 2025a) study the direction of maximum modulation, highlighting its relevance to interpret TI results, but they also use the inefficient maximization-minimization iterative optimization process to compute it.

In this work, we mathematically derive a compact closed-form expression for the direction of maximum modulation depth, 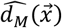, and validate it through simulations. As the discovery of this formula is the major novelty of this work, we show it here in advance:

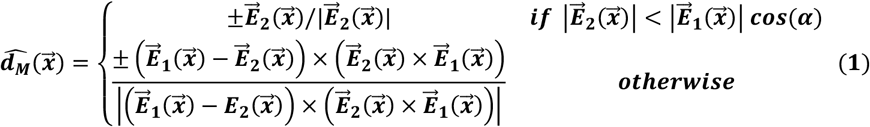

Where 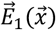 and 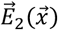 are the amplitudes of the two interfering electric fields and α is the angle between them, considering the same assumptions made by (Grossman et al., 2017) for their closed-form expression of the maximum modulation depth intensity. To our knowledge, there is no analytical expression for this direction published so far, except in one of our conference works (Fernández-Corazza et al., 2019). With a few more steps, we provide a complete mathematical derivation of the expression given by (Grossman et al., 2017) for the modulation depth intensity, given its relevance in the context of TI EBS and the lack of a derivation of it in previous TI EBS works. Finally, we also found that there is another orientation where the modulation depth reaches a local maximum that, in some brain regions, has a similar strength of the maximum modulation depth. As another novelty of this work, we derive closed-form expressions of this secondary maximum modulation depth and of its directions. To our knowledge, this was unnoticed before.

### 2 Modulation depth formulas

In TI, the magnitude of interest is the modulation depth, which indicates how strongly the neurons are being modulated or affected by the stimulation. In TI, a higher modulation depth means stronger potential for neuronal activation. Suppose that 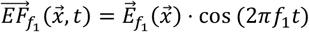 and 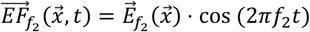 are the two interfering electric fields that oscillate at frequencies *f*_1_ and *f*_2_ = *f*_1_ + Δ*f*, and with amplitudes 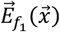 and 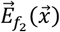, respectively. Note that the amplitudes only depend on the spatial location 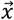 and not on the time *t*. To simplify the formulations, we use the same assumptions made by (Grossman et al., 2017). Without loss of generality, we assume that the two interfering fields are 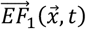 and 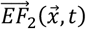 (instead of 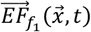 and 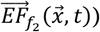) with amplitudes 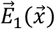 and 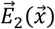 respectively, where 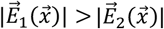, and where the angle α between 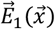 and 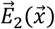 is lower than 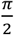. Note that 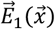 is not everywhere 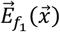 and similarly for 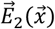 and 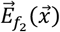 because at a specific position 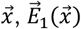 is always the strongest electric field generate by one of the two frequencies *f*_1_ or *f*_2_, and 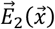 is the weakest electric field generated by the other frequency. Moreover, one of the amplitudes of the two fields 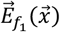 and 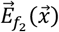 might change its sign when converted to 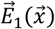 and 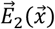 to comply with the assumption made for α.

The modulation depth at a specific unitary direction 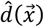 at each point of the space 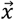, 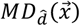, can be calculated as a difference between instructive and destructive interference (Grossman et al., 2017):

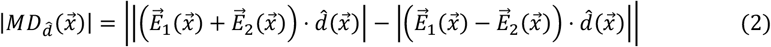

Note that the expression in Eq. (2), as well as all the analysis that follows, does not depend on the time varying electric fields but only on their amplitudes. The formula of Eq. (2) is equivalent to that in (Huang et al., 2020):

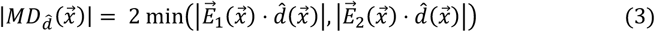

Now, without assuming any specific direction, the formulation for the maximum modulation depth is (Grossman et al., 2017):

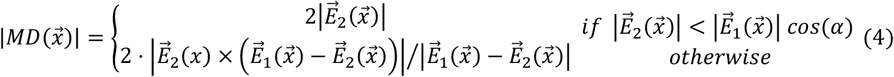

However, along which direction Eq. (4) holds, which we found to be Eq. (1), was not presented before. From now on, we skip the explicit dependency on 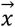 and the upper arrow to indicate vectors to simplify the notation.

## 3 Deduction of maximum modulation depth direction

In Eq. (4) there are two cases: **Case I** occurs in regions of the space where the projection of *E*_1_ onto the direction of *E*_2_ is larger than |*E*_2_| (i.e. |*E*_1_|cos (α) > |*E*_2_|), depicted in Fig. 1(A); and **Case II** holds where the projection of *E*_1_ onto the direction of *E*_2_ is smaller than |*E*_2_| (i.e., |*E*_1_|cos (α) ≤ |*E*_2_|), which is depicted in Fig. 1(B).

**Figure 1.**
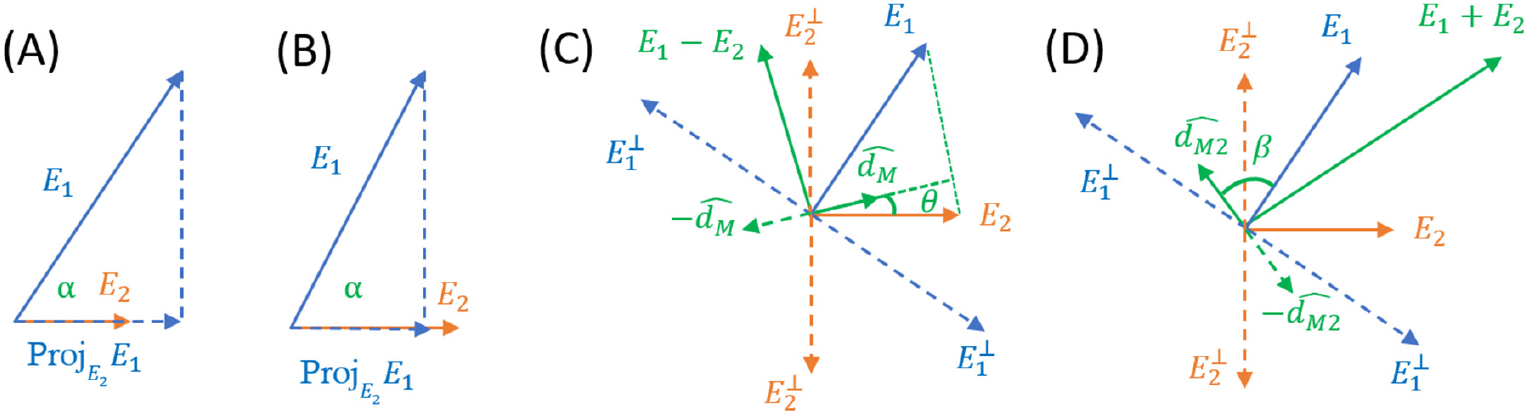
Schematic of Cases I (A) and II (B), where 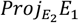 denotes the projection of E_1_ onto the direction of E_2_, and α is the angle between the two vectors. (C) Schematic showing the directions of maximum modulation depth 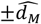 for Case II, which are perpendicular to the E_1_ − E_2_ vector. (D) Schematic showing the directions of second maximum modulation depth 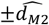,valid for both Cases, which are perpendicular to the E_1_ + E_2_ vector. The directions perpendicular to E_1_ and E_2_, namely 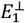 and 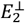, are shown in (C) and (D) as reference because in these directions the modulation depth is zero. Note that in this figure we show the amplitudes of the electric fields as static vectors, without considering the time instance.

For two interfering sinusoidal signals in 3D, the amplitude of the interference at a given unitary direction 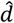 is given by Eq (2). The goal is to find the unitary vector 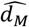 where the maximum of the interference amplitude occurs by analyzing the signs of 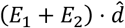 and 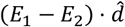 for each Case. Without loss of generality, we also assume that *E*_2_ is located on the *x* axis and that *E*_1_ lies in the first quadrant, as depicted in Fig. 1. We also assume that 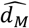 lies in between *E*_1_ and *E*_2_ and that 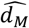 belongs to the plane defined by *E*_1_ and *E*_2_, as demonstrated in (Huang and Parra, 2019).

### 3.1 Case I

In this case, the projection of *E*_1_ onto *E*_2_ is larger than |*E*_2_| (Fig. 1, left). Due to 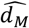 lying between *E*_1_ and *E*_2_, (*E*_1_ + *E*_2_) · 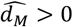. Also, because the projection of *E*_1_ onto *E*_2_ is larger than |*E*_2_|, the projection of *E*_1_ onto any *d* in between *E*_1_ and *E*_2_ is always larger than the projection of *E*_2_ onto 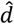 (see Fig. 1). Thus, 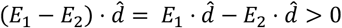.

Then, from Eq. (2):

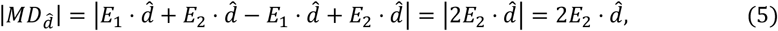

since 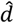 lies between *E*_1_ and *E*_2_ and thus the dot product 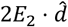 is positive.

If we want to find the unitary direction 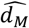 such that 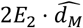 is maximum, the answer is obvious:

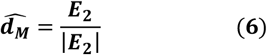

Finally, replacing Eq. (6) in Eq. (5), the maximum modulation depth for Case I is:

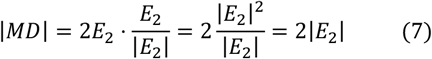

As in (Grossman et al., 2017) formula of Eq. (4), first condition.

### 3.2 Case II

In this case, the projection of *E*_1_ onto the direction of *E*_2_ is smaller than the module of *E*_2_. But still, the module of *E*_1_ is larger than the module of *E*_2_ (Fig. 1, right). In this case, 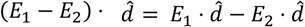 can be either positive, negative or zero. So, we consider three subcases, II-A, II-B, and II-C respectively. We start from the case where 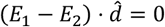, which derives to 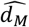. Then, we show that the other two cases can be discarded.

#### 3.2.1 Case II-A

In this case, 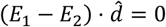, or equivalently 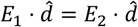, meaning that 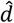 will be the direction in which the projections of both vectors are equal, as shown schematically in Fig. 1(C). It can be observed that 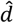 is oriented along the ‘height’ of the triangle with base 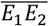. Then, to find 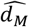 in ℝ^3^ we have a set of conditions:

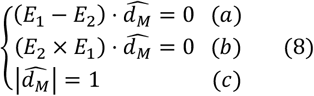

Condition (a) is the one of this subcase II-A, condition (b) means that 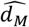 belongs to the plane defined by *E*_1_ and *E*_2_ and condition (c) forces 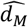 to be unitary. Based on the properties of the cross product: *(i)* α · (*b* × *c*) = *b* · (*c* × α) = *c* · (α × *b*) and *(ii)* 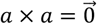, we deducted that *d*_*M*_ = (*E*_1_ − *E*_2_) × (*E*_1_ × *E*_2_) satisfies conditions (a) and (b).

To verify (a), we can write (*E*_1_ − *E*_2_) · *d*_*M*_ = (*E*_1_ − *E*_2_) · ((*E*_1_ − *E*_2_) × (*E*_2_ × *E*_1_)), and using identities *(i)* and *(ii)*, 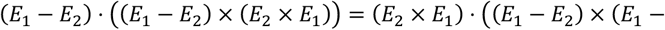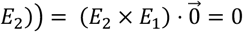.

To verify (b), we can write (*E*_2_ × *E*_1_) · *d*_*M*_ = (*E*_2_ × *E*_1_) · ((*E*_1_ − *E*_2_) × (*E*_2_ × *E*_1_)), and using identities *(i)* and *(ii)*, 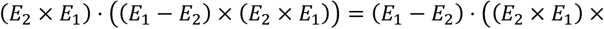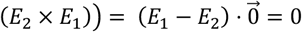.

To satisfy (c), we just normalize *d*_*M*_ obtaining the 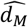 expression of Eq. (1), repeated here for clarity:

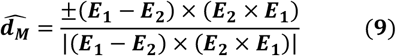

Eq. (9) shows the explicit formula of the direction where the maximum modulation depth of the interfering signals occurs, and it constitutes the major novelty of this work. The ± sign is because as everything is alternate current (AC), there are in fact two opposite directions where the intermodulation depth is maximum. Strictly speaking, one can refer to only one orientation of maximum modulation depth. Note that formula of Eq. (9) is not given in (Grossman et al., 2017).

We show next that Cases II-B and II-C lead lower modulation depths and thus, there is a maximum at 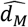.

#### 3.2.2 Case II-B

In this subcase, 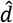 is in between 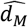 and *E*_1_, thus 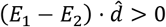 (see Fig. 1c). From Eq. (2),

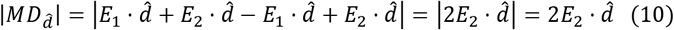

The last equality holds because from Fig. 2, 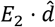 is always positive. But as 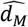 is closer to *E*_2_ than 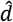 then 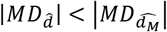.

#### 3.2.3 Case II-C

In this subcase, 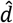 is in between 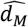 and *E*_2_, thus 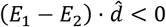 (see Fig. 1c). From Eq. (2),

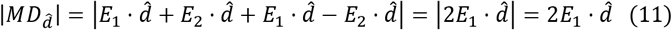

The last equality holds because from Fig. 2, 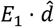 is always positive. But as 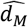 is closer to *E*_1_ than 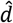 then 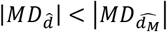.

In summary, there is a maximum at 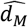 (and at 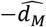 because everything is AC). However, there are secondary maximums in the second and fourth quadrants, as shown in Section 4.

#### 3.2.4 Derivation of Grossman et. al. formula for Case II

Here we derive Grossman et al (2017) closed form expression for the maximum modulation depth of Case II, i.e., the bottom row of Eq. (4). Using the condition that we found 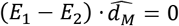 and consequently 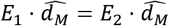, the maximum modulation depth from Eq. (2) can be written as:

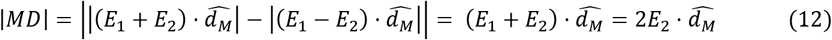

Using the dot product property, 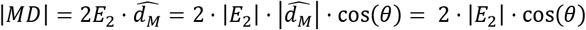, where *θ* is the angle between *E*_2_ and 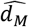 (see Fig. 1c), and 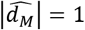 because it is unitary by definition. But 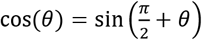 (a trigonometric property), and 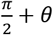 is the angle between *E*_1_ − *E*_2_ and *E*_2_ (see Fig. 2). Remember that in this subcase II-A 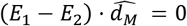, meaning that *E*_1_ − *E*_2_ is perpendicular to 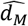 (see Fig. 2). Then:

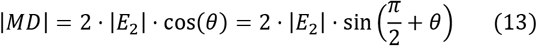

Now, using the cross-product property |α × *b*| = |α||*b*||sin(α)|, and because 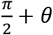 is the angle between (*E*_1_ − *E*_2_) and *E*_2_:

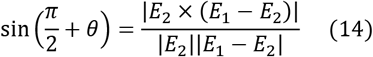

Remember that 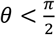, thus 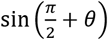 is always positive.

Finally, replacing Eq. (14) in Eq. (13), we get

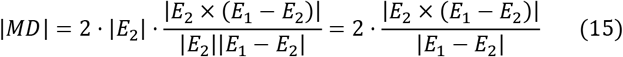

Which is (Grossman et al., 2017) formula for Case II.

## 4 Direction of secondary modulation depth maximum

It is obvious that there are four nulls in the directions perpendicular to *E*_1_ and to *E*_2_, namely 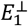 and 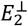 respectively. These directions are shown in Fig. 1c and Fig. 1d. And because the modulation depth is always positive, there should be another two maximums between these two orientations. We found that these secondary maximums are located at the orientation perpendicular to (*E*_1_ + *E*_2_), i.e. *d* · (*E*_1_ + *E*_2_) = 0. We also found the closed-form expression of these directions, valid for both Cases I and II (see Fig. 1d):

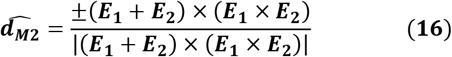

Note that Eq. (16) is analogous to Eq. (9). Following similar steps of sections 3.2.2 and 3.2.3, considering first a direction *d* in between 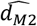 and 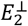 (i.e. *d* · (*E*_1_ + *E*_2_) > 0), and then a direction 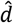 in between 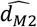 and 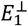 (i.e. *d* · (*E*_1_ + *E*_2_) < 0) it is possible to demonstrate that there is a local maximum of the modulation depth at 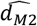. In fact, these proofs demonstrate that there are only four maximums located at 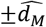 and at 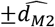 in standard TI stimulation with two sinusoidal frequencies, if the two electric fields are not exactly aligned. Thus, there are no more maximums besides the ones we found. If *E*_1_ and *E*_2_ are exactly aligned, unlikely to occur in a real case scenario, there will be only two maximums at 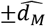.

Moreover, following similar steps of section 3.2.4 to get Eq. (15) from Eq. (12), it is possible to obtain the modulation depth at this secondary orientation, which turns out to be:

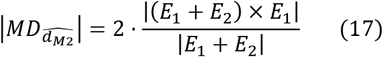

In this case, the trigonometrical identity 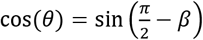 is used instead of 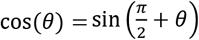, where β is the angle between 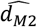 and *E*_1_ (see Fig. 1d). Note that Eq. (17) is analogous to Eq. (4) – Case II, same as Eq. (15), but for the secondary maximum it is valid in both Cases I and II.

## 5 Validation using simulations

We validated the theoretical findings by means of computational simulations of TI TES on a finite element realistic head model with seven tissues (scalp, compact skull, marrow bone, cerebrospinal fluid, eyeballs, gray matter and white matter), ∼6.5 million tetrahedra (∼800 k belonging to the brain) and ∼1.1 million nodes. We assigned literature conductivity values for each tissue as follows: 0.2S/m for the WM (Koessler et al., 2017), 0.006 S/m for the skull (Fernandez Corazza et al., 2018), 0.3 S/m for the scalp (Fernandez Corazza et al., 2018), 1.6S/m for the CSF (Chauhan et al., 2018), 0.45 for the GM (McCann et al., 2019) and 1.55 S/m for the eyeballs (Lindenblatt and Silny, 2001). We reused a realistic head model for electrical fields in the quasistatic approximation from a previous work, where the construction process is highly detailed (Fernandez Corazza et al., 2018).The four electrode positions for applying the two interfering patterns of 1mA each were selected to mimic the current injection pattern used in (Violante et al., 2023) for real experiments (Figs. 3a and 3b). Computations were done separately for each frequency *f*_1_ and *f*_2_. As a result, two independent static vector field amplitudes 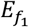 and 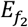 were obtained in each tetrahedral element of the 3D head model and sorted out to form *E*_1_ and *E*_2_ following the assumptions described in section 2. Those static vector amplitudes due to the linearity of the Poisson equation and constant tissue conductivities represent the amplitudes of the interfering fields. Fig. 3c shows a sagittal slice at approximately the same height as the electrode locations with the regions of Case I (blue) and the regions of Case II (green).

**Figure 3.**
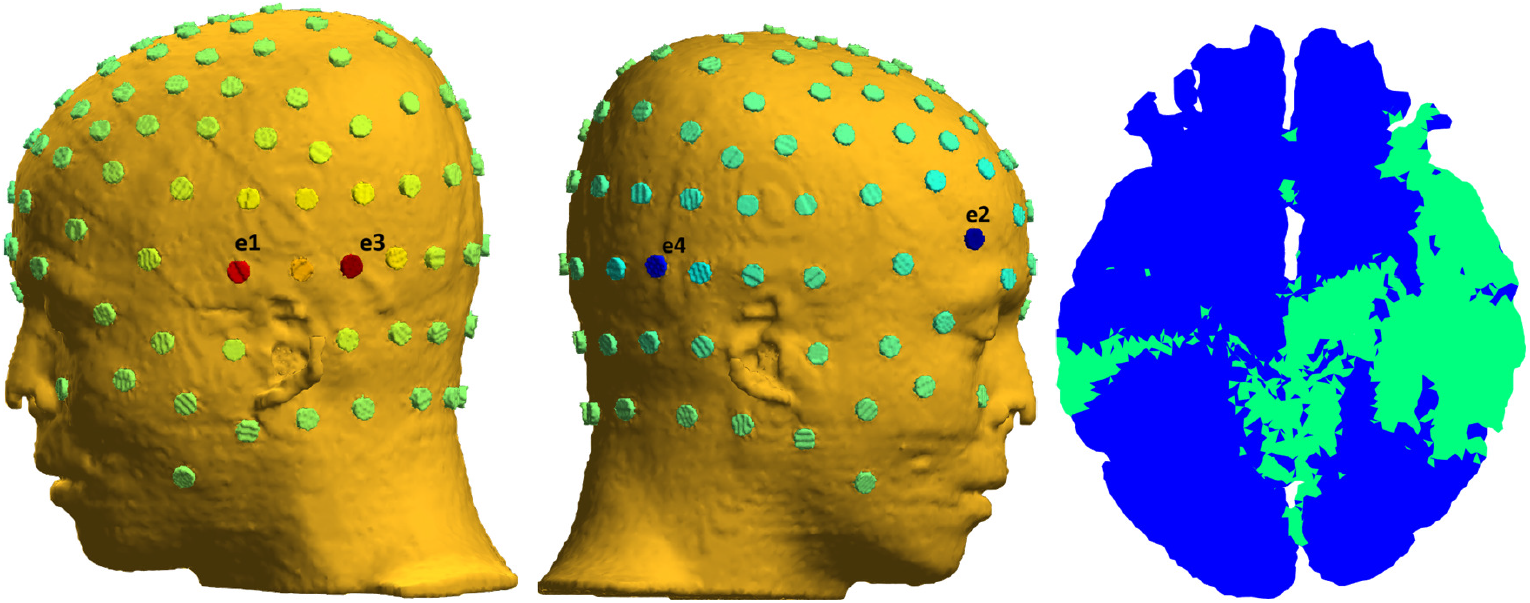
(a) Left view with electrodes positions e1 and e3. (b) Right view with electrodes positions e2 and e4. The current injection pairs are e1-e2 for one frequency and e3-e4 for the other. (c) Sagittal slice of the brain (white matter and gray matter) at the approximate height of the electrode positions showing the regions where Cases I (blue) and II (green) hold.

For each tetrahedral element of the brain, we computed the modulation depth intensity in three different ways. First, we computed the maximum modulation depth direction of Eq. (1) and used it as 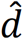 in Eq. (2) to compute the maximum modulation depth (Fig. 4A). Second, we computed the modulation depth directly using (Grossman et al., 2017) formula of Eq. (4) (Fig. 4B). Third, we generated two sinusoidal waveforms at 1000Hz and at 1010Hz (sampling rate of 40KHz and 1 second of duration), multiplied each of them by the two field amplitudes 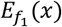 and 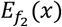 respectively obtaining the dynamic fields 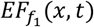 and 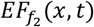, computed the sum of the electric fields at each tetrahedral element of the brain and at each time point, projected the fields onto the optimal direction of Eq. (1) and used MATLAB’s “envelope” function to obtain the modulation depth of the envelope of the time varying signals (Fig. 4C). It can be observed that the three ways of computing the maximum modulation depth give identical results, thus verifying that the direction of maximum modulation depth of Eq. (1) is correct. Note that for the first two methods (Figs. 4A and 4B), the computation does not require simulations of the raw signals evolving with time. We also computed the modulation depth along the secondary direction of Eq. (16) following the same three procedures, where in the second procedure we used Eq. (17) which is the analogue to (Grossman et al., 2017) formula of Eq. (4). In Fig. 4D we show only one image because again all three procedures for computing this map are equal. At the regions near points “b” and “c”, the intensities of the main (Fig 4C) and the secondary (Fig 4D) maximums are almost the same, while in the region near point “a” at the left hemisphere, the main maximum dominates over the secondary one.

**Figure 4.**
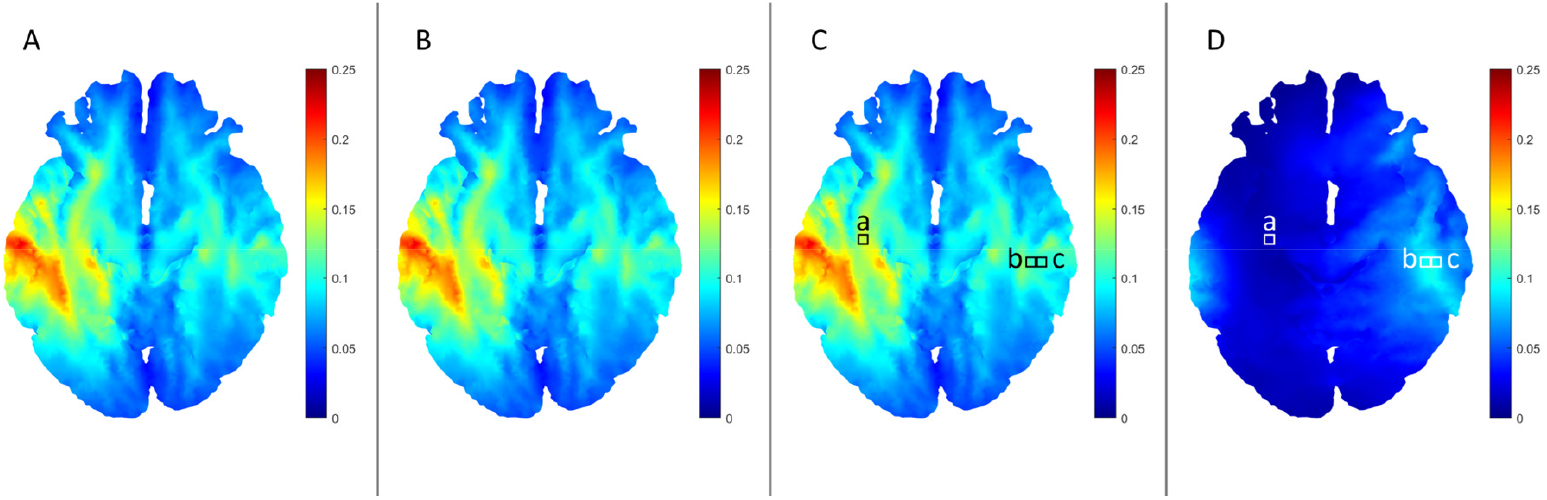
Sagittal slices of the brain (white matter and gray matter) showing the maximum modulation depth (in [V/m]) computed (A) using the novel closed-form expression of the maximum modulation depth direction 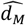 of Eq. (1) inserted into the general formula of Eq. (2), (B) using Grossman et al formula of Eq. (4), and (C) generating the two raw interfering electric field waveforms projected onto the direction of Eq. (1) and computing the depth using MATLAB’s “envelope” function. Subplot (D) shows the secondary maximum modulation depth at the same scale computed using the closed form expression of Eq. (16) inserted into Eq. (2), likewise (A). We also computed the secondary modulation depth using Eq. (17), likewise (B), and using the waveforms, likewise (C), yielding in both cases the exact same results as (D) (not shown). In subplots (C) and (D) we highlight three positions “a”, “b” and “c” used as representative examples.

In Fig. 5 we show, for the same slice of the brain, the orientations of the two electric field amplitudes *E*_1_ and *E*_2_, plus the orientations of the main and secondary maximum modulation depths. It can be observed that in the Case I regions, the maximum modulation depth orientation matches one of the electric field directions (for instance, position “a” in Fig 5). In the Case II regions, the maximum modulation depth orientation is different than the two interfering electric fields. One interesting phenomenon can be observed in the zoomed region of Fig. 5, where the maximum modulation depth orientation changes abruptly, as it occurs between positions “b” and “c”. It can be observed that in those cases, the main and secondary maximum modulation depth orientations are like 90 degrees apart (but not exactly) and switch positions. When the two electric fields are almost perpendicular and of similar strength, the orientation of maximum modulation depth is approximately at 45 degrees of both fields, but the direction of secondary maximum depth has also a large modulation depth. Thus, when a slight change in the angle of the fields occurs, the previous secondary direction 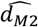 now takes the lead replacing 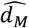 as the one with maximum modulation depth. Overall, the switching phenomenon is smooth, but the maximum modulation depth orientation can have abrupt jumps. To illustrate this phenomenon, we plot in Fig. 6 the waveforms projected onto the orientations of maximum modulation depth 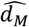 and 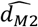, for the three representative positions “a”, “b” and “c” of Fig. 5.

**Figure 5.**
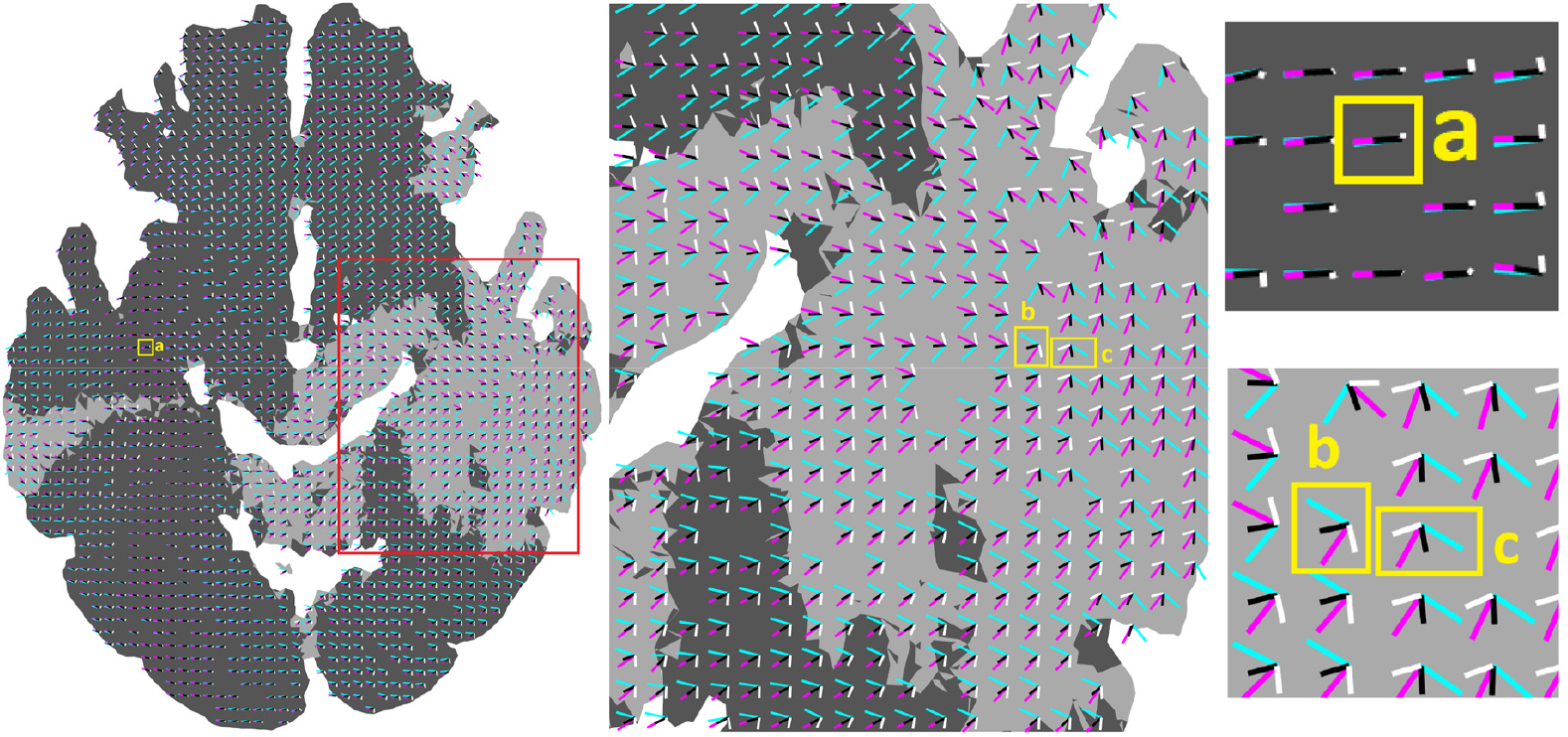
Electric field orientations (E_1_ in cyan and E_2_ in magenta), orientation of maximum modulation depth 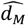 (in black) and of secondary maximum 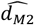 (white) for the same sagittal brain slice of Fig. 4, distinguishing where Cases I (dark gray) and II (light gray) hold (left). A zoomed version of the red rectangle region is shown in the middle and regions near the points of interest are shown on the right with an even larger zoom. In the Case I regions (dark gray), 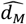 matches one of the electric fields. Note how 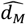 switches to 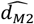 when moving from “b” to “c” and vice versa. The time waveforms of representative positions “a”, “b” and “c” are shown in Fig. 6. In all cases the 3D vectors are normalized and only the x and y components are depicted. 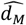 and 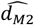 lines were shortened a little bit more to improve clarity when they overlap with one electric field (such as in position “a”).

**Figure 6.**
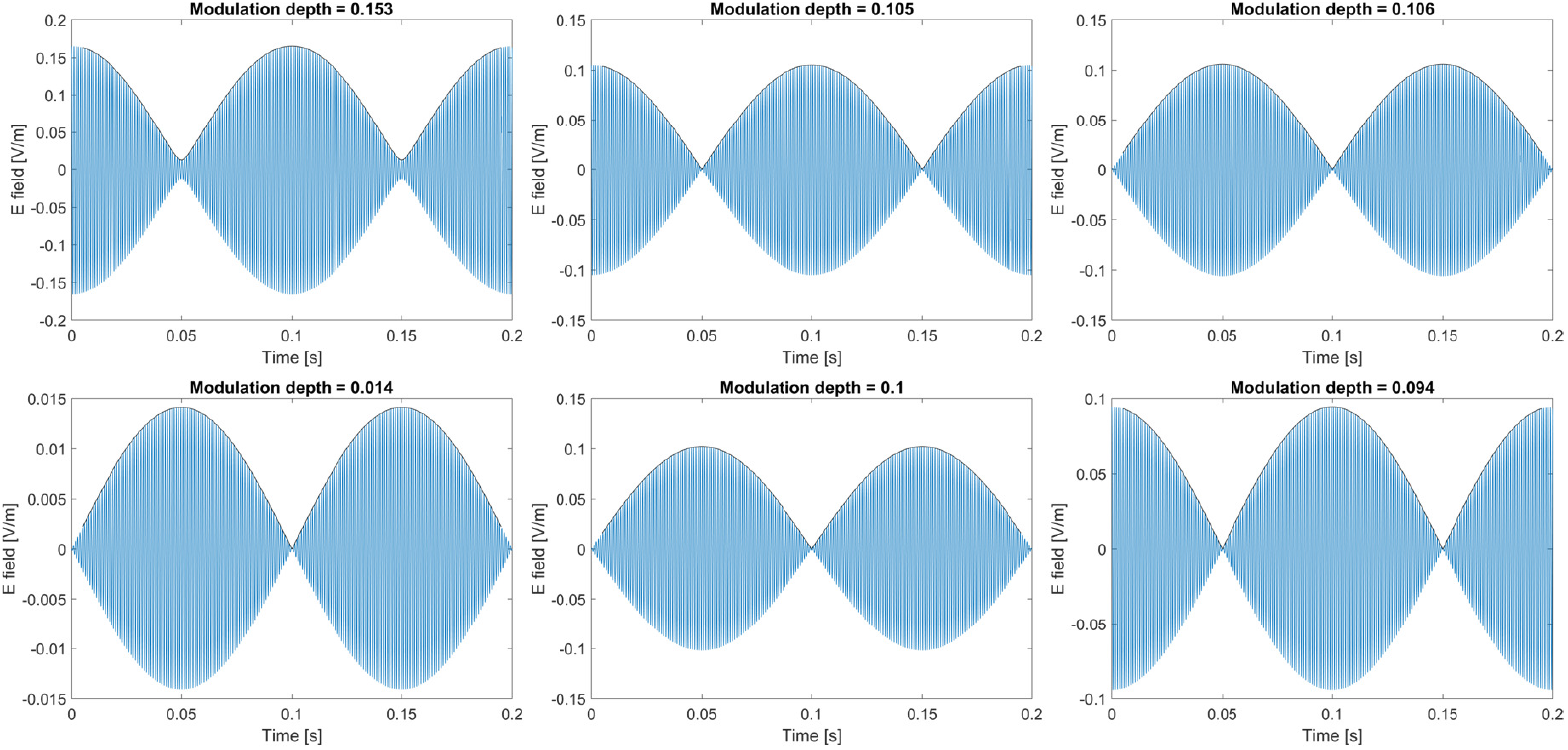
Waveforms (blue) and envelopes (black) for three representative examples at the three locations “a” (first column), “b” (second column) and “c” (third column) of Fig. 5 for one period of the beating amplitude variation in time (200ms = 1/5Hz) or two periods of the envelope frequency (100ms = 1/10Hz). The top row shows the waveforms projected onto the orientation of maximum modulation depth 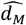 and the bottom row shows the waveforms projected onto 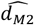. The numerical value of the modulation depth is shown in the title of each subplot. In the first column, example “a”, Case I holds, and the two electric fields are almost aligned. Thus, 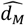 matches one of the electric fields and 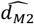 is much smaller. In the second and third examples, both electric fields are almost perpendicular and thus, the modulation depths are high for both 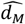 and 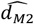.

In Fig. 6 it is possible to observe that in all cases the envelopes at 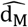 and 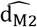 are in anti-phase (90 degrees difference in time). This means that, at one timepoint, the peak of the beating low-frequency signal can be at maximum along the primary direction and minimum along the secondary direction but swap values in half of the envelope frequency (50ms in Fig. 6). Note that the result of summing two signals at two different frequencies results in a carrier wave with average frequency (*f*_1_ + *f*_2_)/2 (1005Hz in our simulations) that has an amplitude modulated with a frequency |*f*_1_ − *f*_2_|/2 (5Hz). However, taking the envelope involves a rectification process where the intensity is the square of the amplitude, resulting in an envelope of twice the frequency, i.e. 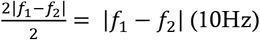.

We also performed a brute force analysis where we computed for all elements of the brain the modulation depths of the envelope of the sum of the two interfering electric fields, projected onto ∼2500 fixed directions uniformly covering the entire 4 π solid angles. We compared the modulation depths with the modulation depth of the signals projected onto 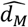. In all cases, as expected, the 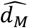 resulted in the largest modulation depth among all tested directions. This serves as an empirical validation of the direction of maximum modulation depth of Eq. (1) and of the assumptions described at the end of section 2.

In Fig. 7 we show the modulation depth for all the angles of the plane defined by *E*_1_ and *E*_2_ (top row) and for all the 3D directions (bottom row), at the three positions “a”, “b” and “c” of Fig. 5. The directions 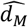 and 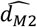 are also plotted and once again, we validate that they correspond to the peaks of the modulation depth. It can be observed that besides 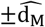 and 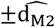 there are no other maximums. It is interesting to observe how the modulation depth is broader in case “a” and more focal in terms of direction in cases “b” and “c”, where 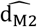 is almost as strong as 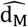. Note also how the modulation depth smoothly diminishes outside the plane defined by *E*_1_ and *E*_2_.

**Figure 7.**
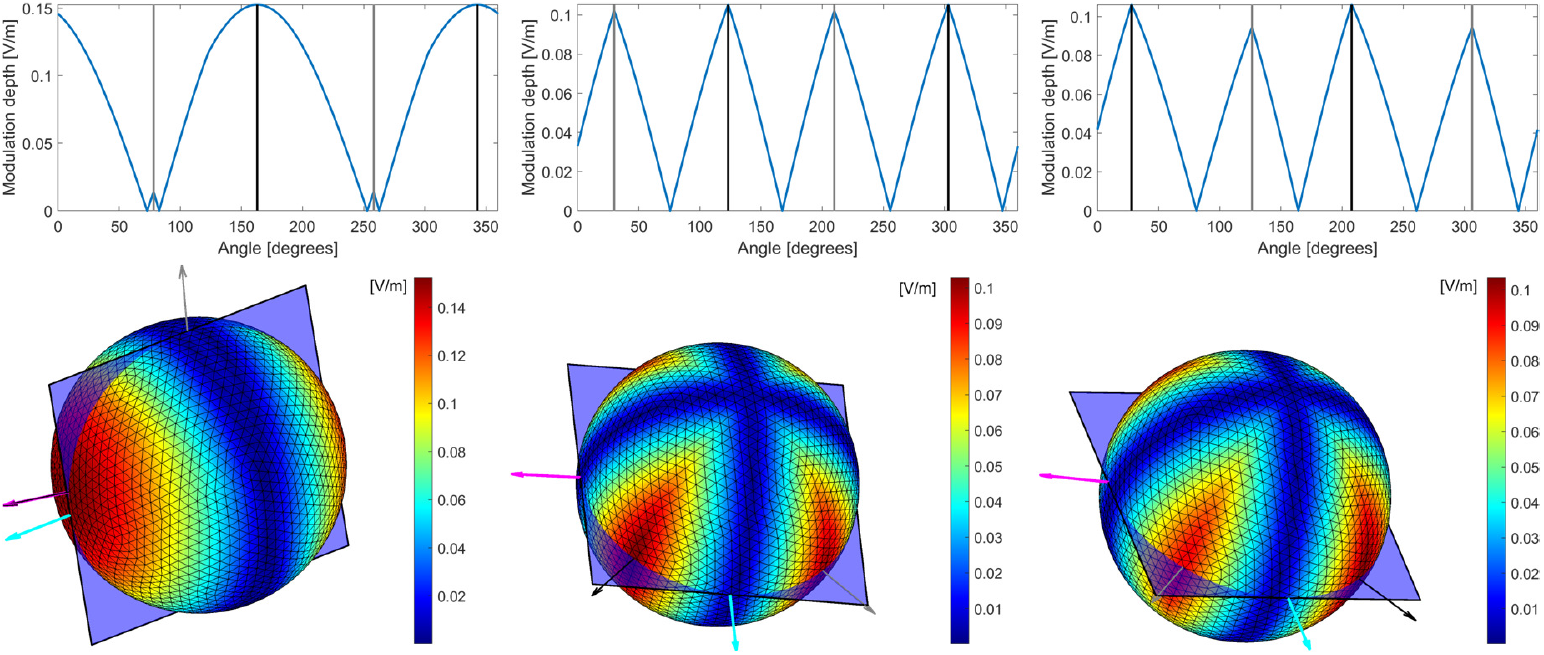
Examples of the brute force modulation depth calculations for positions “a” (first column), “b” (second column), and “c” (third column) of Fig. 5. The top row shows the modulation depth in the plane defined by E_1_ and E_2_ as a function of the angle, where directions 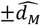 and 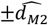 are indicated with black and gray vertical lines respectively. The bottom row shows the modulation depths along all directions of a 3D sphere encoded in color. The directions of the E_1_ and E_2_ fields are shown as cyan and magenta arrows as well as 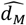 and 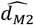 as black and gray arrows respectively. The plane defined by E_1_ and E_2_ is also shown.

In previous studies it was shown that by changing the current intensity ratio between the two electrode pairs, it was possible to spatially steer the spot of maximum modulation depth slightly to one side or to another (Fernández-Corazza et al., 2019; Grossman et al., 2017). Here we show that the directions of the main and secondary maxima can also be slightly steered to one side or to the other using the same technique, as in the two examples depicted Fig. 8. In one example, we applied 1.2mA in the e1-e2 pair and 0.8mA in e3-e4 pair, and in the other we applied 0.8mA and 1.2mA in the e1-e2 and e3-e4 pairs respectively. Note that in Case I the steering might not be strong enough to modify the direction of maximum modulation depth because in this case it matches one of the electric fields. However, the secondary maximum field does steer (see zoomed region in Fig. 8, top row first column). In Case II, the steering occurs in the directions of both the main and secondary maximum modulation depth because these directions do not match any of the interfering field directions. Overall, the current injection ratio steering can be useful to easily perform a fine adjustment of either the primary or secondary maximum modulation depth directions to better align any of them to some desired direction, such as the direction of the neuronal columns.

**Figure 8.**
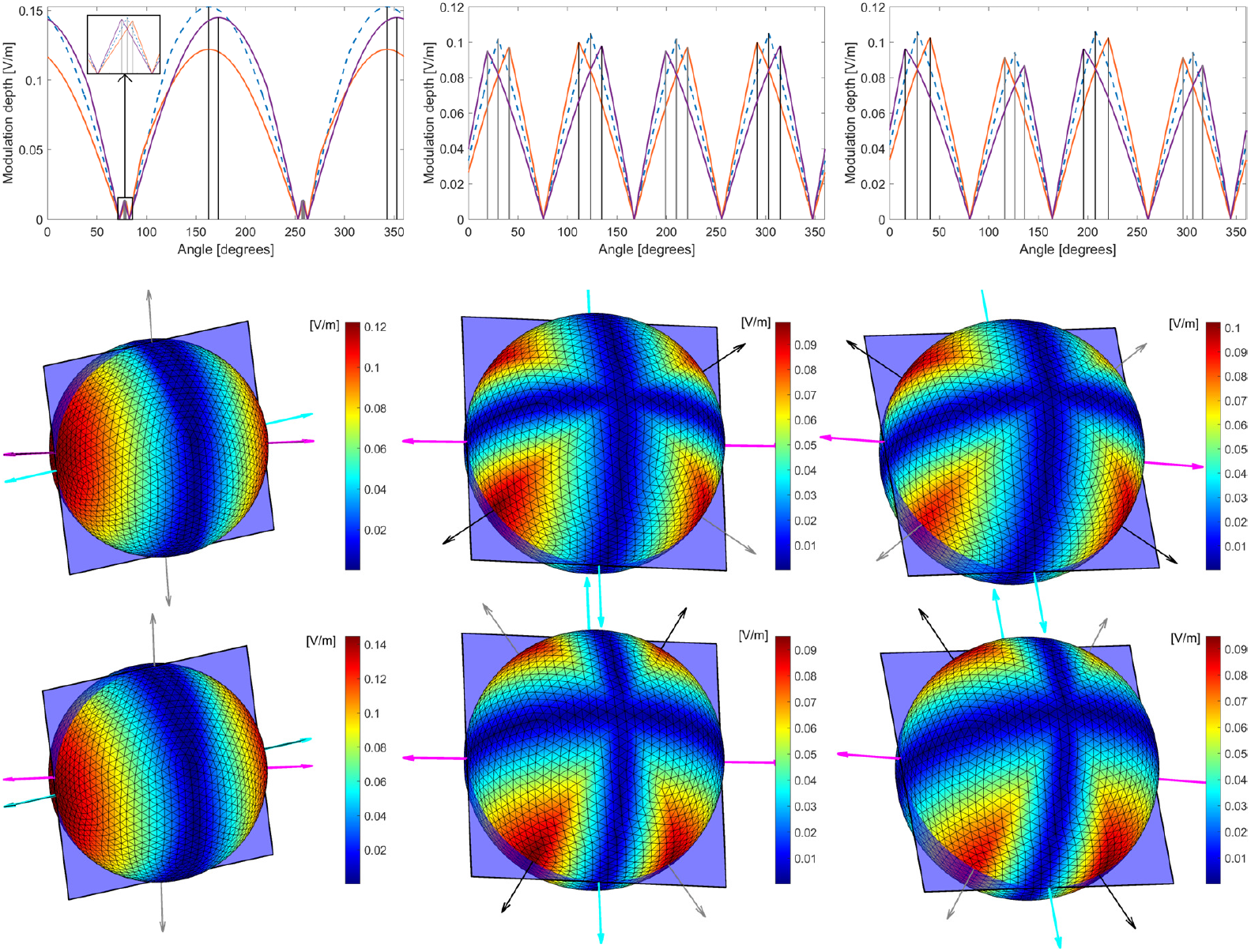
Examples of the directional steering when changing the ratio of the applied current to each frequency pair, for positions “a” (first column), “b” (second column), and “c” (third column) of Fig. 5. The top row shows the modulation depths in the plane defined by E_1_ and E_2_ as a function of the angle for: 1.2mA in the e1-e2 pair and 0.8mA in e3-e4 pair (orange line); 0.8mA in the e1-e2 pair and 1.2mA in e3-e4 pair (purple line); and equal current injection for reference (dashed-blue line, same as Fig. 7 top row). Directions 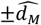 and 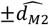 are indicated with black and gray vertical lines respectively. The middle row shows the modulation depths along all directions of a 3D sphere when 1.2mA and 0.8mA are applied to the e1-e2 and e3-e4 pairs respectively and the bottom row shows the modulation depth when the applied currents are 0.8mA and 1.2mA respectively. The directions of the ±E_1_ and ±E_2_ fields are shown as cyan and magenta arrows as well as 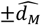 and 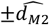 as black and gray arrows respectively. The plane defined by E_1_ and E_2_ is also shown. Note that the scales are different than the scales of Fig. 7, bottom row (equal current injection).

## 6 Comparison with other studies

Other studies state the maximum modulation depth as a maximization-minimization problem that can be solved numerically with iterative solvers (Huang and Parra, 2019; Meng et al., 2025b), based on Eq. (3).

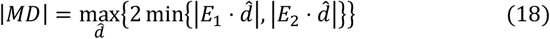

We showed in Section 3 that Eq. (4) is the closed-form solution to Eq. (18). Eq. (4) is preferred over Eq. (18) to avoid time consuming and error prone iterative solvers.

Regarding closed-form expressions, (Wang et al., 2022) propose a formulation that looks similar to Eq. (4) although it does not exactly match:

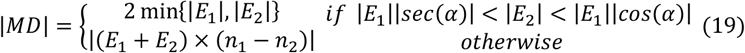

In Eq. (19), 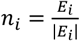 and it is not assumed that |*E*_1_| > |*E*_2_|. First, the condition |*E*_1_||*sec*(*α*)| < |*E*_2_| < |*E*_1_||cos(*α*)| is at least misleading because it impossible to satisfy both conditions simultaneously, since |*E*_1_||*sec*(*α*)| < |*E*_1_||cos(*α*)| or equivalently |*E*_1_| < |*E*_1_||cos(*α*)|^2^ will never occur (|cos(*α*)| < 1 always). To match Eq. (4), if min{|*E*_1_|, |*E*_2_|} = |*E*_2_|, only the second inequality |*E*_2_| < |*E*_1_||cos(*α*)| should hold (same condition of Eq. (4)), and if min{|*E*_1_|, |*E*_2_|} = |*E*_2_|, only the first inequality |*E*_1_||*sec*(*α*)| < |*E*_2_| should hold. Further, the bottom-line expression in Eq. (19) should be

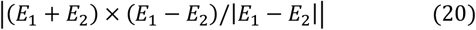

instead of |(*E*_1_ + *E*_2_) × (*n*_1_ − *n*_2_)| to match the expression of Eq. (4). Adding and subtracting 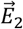 to the first term of Eq. (20) we get | (*E*_1_ + *E*_2_ + *E*_2_ − *E*_2_) × (*E*_1_ − *E*_2_)/|*E*_1_ − *E*_2_|| | (2*E*_2_ + (*E*_1_ − *E*_2_)) × (*E*_1_ − *E*_2_)/|*E*_1_ − *E*_2_| = |2*E*_2_ × (*E*_1_ − *E*_2_)/|*E*_1_ − *E*_2_|| because 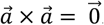. We also checked numerically that in fact both expressions from Eq. (4) and (19) are different. Thus we consider valid only the expression of Eq. (4) (Grossman et al., 2017).

Regarding the direction of maximum modulation depth 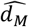, only (Meng et al., 2025b) claim they find it, but they do it by solving the numerical formulation of Eq. (18) and not finding a closed-form expression as we show in this work. They claim they use these additional quations to find 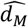:

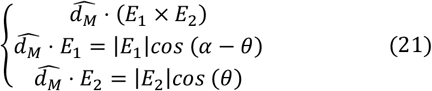

Where 0 is the angle between 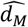 and *E*_2_. However, the expressions in Eq. (21) hold for any 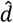 within the plane defined by *E*_1_ and *E*_2_. Thus, expressions of Eq. (21) do not provide additional information to define a unique 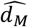 in contrast to the conditions presented in in this work in Eq. (8) that do define a unique 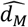.

Moreover, no other study explicitly noted neither existence nor orientation of the secondary maximum modulation depth 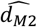, which we provide its closed-form expression in Eq. (17) and a closed-form expression of its modulation depth in Eq. (18).

## Conclusion

In this work we present closed-form expressions for the main and secondary directions of the maximum modulation depth in temporal interference brain stimulation. The direction of maximum modulation depth was not discussed in previous studies that present closed-form expressions for the maximum modulation depth (Grossman et al., 2017; Wang et al., 2022), whereas the existence and direction of the secondary maximum modulation depth was completely overlooked due to the vectorial 3D nature of temporal interference. A use of the closed form expressions presented in this work results in much faster and precise computations than using numerical iterative optimizations as done in recent studies (Huang and Parra, 2019; Meng et al., 2025b). The closed-form expressions found for two possible maxima of comparable strength and anti-phase in time provide new insights and are highly relevant for better understanding and interpretation of both synthetic and real experimental results of TI stimulation by comparing these directions with the neuronal alignment at different locations of the brain (Meng et al., 2025b). We also show that by changing the ratio of the applied currents, it is possible to steer these directions for better alignment. As secondary results, we provide a full derivation of (Grossman et al., 2017) formulation which was missing in the literature and a corrected version of (Wang et al., 2022) formulation. Finally, our analytic 3D results can be generalized beyond neuromodulation to other fields of research in physics, like radar and sensing, signal processing, radiofrequency and optical communications, acoustics and sound engineering where the notion of TI is used.

## Acknowledgements

This work was supported by the ANPCyT PICT 2019-0701, UNLP 11/EI-003, CONICET PIP 11220200101515CO. MFC wants to thank PhD. S. Collavini, PhD. D. Dellavale and PhD. E. Urdapilleta for their helpful input on specific aspects of this paper.

